# Ecology and Function of Bacterial Lipases within the Gut Microbiome

**DOI:** 10.1101/2020.09.08.287425

**Authors:** Thomas C. A. Hitch, Johannes M. Masson, Theresa Streidl, Thomas Fischöder, Ditte Hobbs-Grimmer, Debjyoti Ghosh, Lothar Elling, Nico Jehmlich, Martin von Bergen, Lesley Hoyles, Thomas Clavel

## Abstract

Triglycerides are a major dietary component and excessive intake has been linked to many health conditions. While the impact of host lipases on fat degradation and absorption have been well documented, bacterial lipases have been largely ignored. Here, we identified the landscape of microbial lipases within the gut of mice, pigs, and humans. In total, 373 microbial lipases were identified and formed two major clusters based on protein sequences, with the sequences containing highly conserved motifs. Metagenomic and cultivation approaches identified lipase-positive bacteria as diverse, occurring within seven phyla, and abundant within the human gut (9.1% ± 4.6%). Pathway analysis in lipase-positive species identified a low prevalence of fatty acid utilisation and a higher frequency of glycerol metabolism. In vitro testing with stable isotopes confirmed that human gut bacteria can utilise the released glycerol. Overall, this study highlights the role of bacterial lipolytic activity in triglycerides metabolism within the gut.

## Introduction

The gut microbiota is a complex ecosystem that consists of hundreds of microbial species collectively producing millions of proteins [1–5]. Mechanistic understanding of their role in health and disease is made difficult due to this complexity, leading to the use of synthetic communities [6, 7]. Even when the complexity of the microbiota is reduced, mechanistic understanding is hampered by the lack of meaningful annotations for a majority of microbial functions within the gut [8]. One approach for improving functional annotations is the targeted analysis of specific functions, as previously conducted for β-glucuronidases within both the human and murine gut [9, 10] and also for bile acid metabolism [11].

The prevailing knowledge about fat absorption is that it occurs within the small intestine, specifically in the proximal region [12, 13], according to studies conducted in the 1950s [12, 14, 15], 1960s [16] and 1970s [13, 17–19]. However, in light of findings which suggest fat absorption differs greatly depending on fat amount [20] and type provided [21], as well as the potential role of the gut microbiome [22], these assumptions should be re-evaluated and confirmed to provide comprehensive overview of mammalian fat absorption. To date, the role of bacterial lipases on fat degradation and host metabolism has only been investigated a handful of times. Oral administration of a transgenic lipase-producing bacterium was shown to significantly increase fat absorption and improved steatorrhea in pigs that had undergone pancreatic ligation [23]. Carp (*Cyprinus carpio*) fed with LipG1, a bacterial lipase isolated from the gut of carp, significantly increased lipolytic activity and weight gain when fed palm oil, a cheap and industrially advantageous feed [24]. Whilst these studies highlighting both, the health and economic benefits to better understand bacterial lipases and fat degradation, bacterial lipases within the gut are currently understudied compared to host-derived lipases [25–27].

In this paper, we studied bacterial lipases present within the gut using a combination of metagenomics, bacterial cultivation, and biochemical methods. This includes identification of the diversity of lipase enzymes, taxonomic diversity of lipase-positive species, and metabolic implications for them.

## Results

### Specific bacterial lipase repertoire within the mammalian gut

While functionally diverse, lipases and esterases share structural and sequence similarity as they are members of the protein family of alpha/beta hydrolases [28]. To ensure correct identification of lipases within gut microbiota gene catalogues (human, mouse, pig), annotation was conducted against three reference databases. False-positive identification was prevented by selecting proteins with matches in two of the three databases. Multiple conserved structural features exist within lipases, including the alpha-beta fold and motifs such as GxSxG at the active site [29]. Hence filtering was conducted based on the requirement that proteins must contain this motif. This reduced the number of total matches by >45% (**Table 1**).

**Table 1:** Annotation of gut bacterial protein catalogues for lipases. The table indicates the total number of proteins within each host catalogue, in million, followed by the number identified as being lipases throughout the different steps of the workflow.

| *Database* | **Human (9.9 million) [2]** | **Mouse (4.5 million) [4]** | **Pig (7.7 million) [3]** |
| --- | --- | --- | --- |
| *LED* | 10,988 | 5,658 | 6,490 |
| *EC* | 14 | 5 | 20 |
| *EGGNOG* | 239 | 274 | 192 |
| **Annotation in ≥2 databases** | 175 | 250 | 158 |
| **GxSxG motif** | 116 | 178 | 79 |

The identified gut lipases were placed into the latest classifcation system for bacterial lipases [30] based on sequence identity using an all-against-all annotation approach (**Figure 1a**). While some gut bacterial lipases were related to family 1.1, 1.2, 1.3, 1.4 and 7, the majority grouped together to form two loosely connected clusters, labelled as Cluster 1 and 2. These two clusters contained lipases originating from the microbiota of humans, mice, and pigs, while some smaller clusters were host-specific.

**Figure 1:**
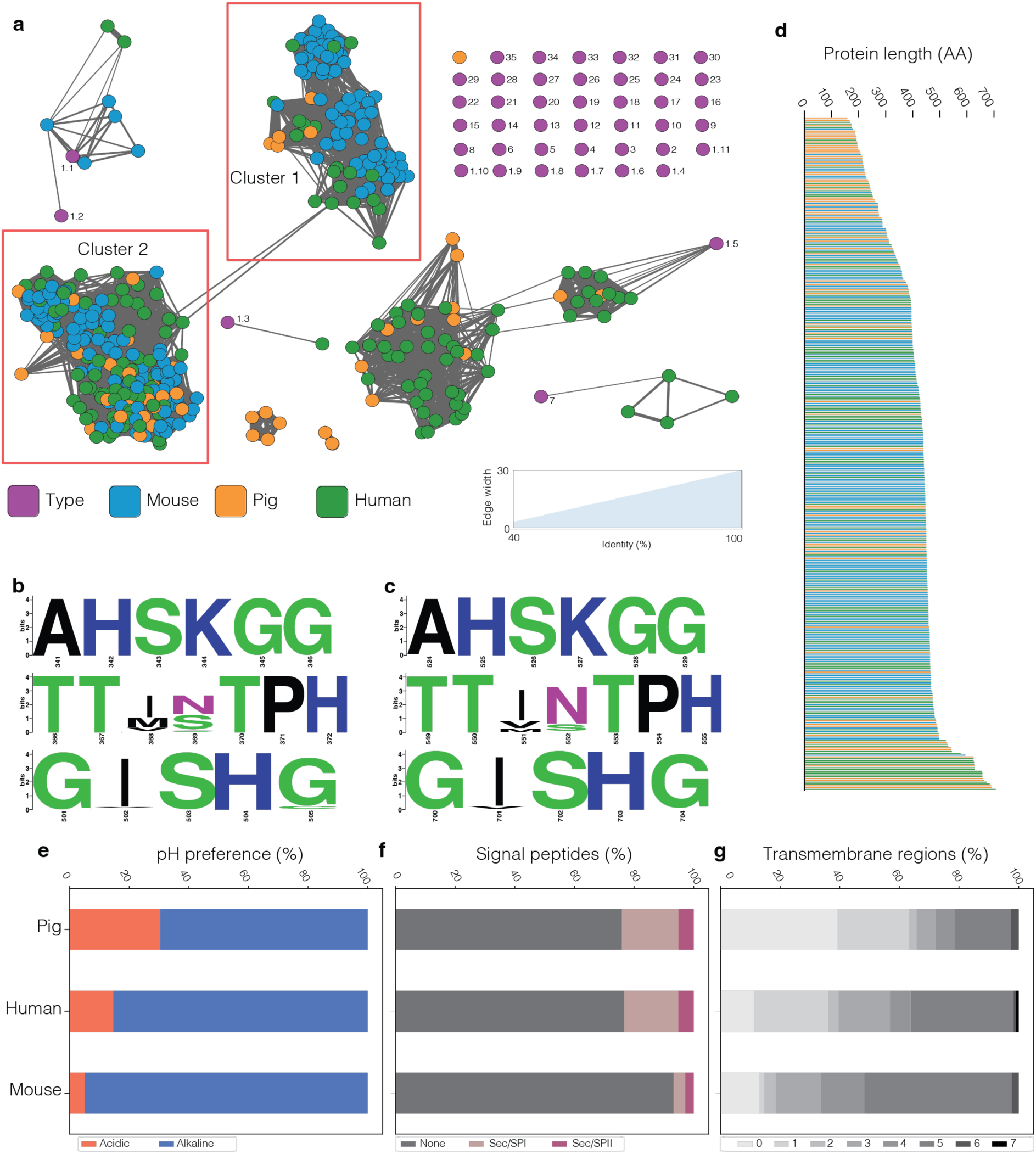
Protein sequence analysis of gut bacterial lipases. **a.** Network analysis of sequence identity shared between the identified bacterial lipases. Each node represents a single lipase coloured based on its host of origin. Edge lengths and widths are proportional to the sequence identity shared between the two nodes. Cluster 1 and 2 are highlighted with red boxes. **b - c.** The conserved motifs identified within Cluster 1 (**b**) and Cluster 2 (**c**) along with their respective positions in the alignment and coloured based on amino acid features. **d**. Protein length variation across the lipases, coloured based on their host of origin. **e - g.** The features of these enzymes were predicted using AcalPred [31] for determining if enzymes were alkaline or acidic (**e**), SignalP [32] for prediction of signal peptide sequences (**f**) and TMHMM [33] for prediction of transmembrane regions (**g**).

Due to the high sequence identity observed between proteins within Cluster 1 and 2, further sequence analysis was conducted to identify conserved amino acid motifs. Within Cluster 1, the motifs AHSKGG and TTxxTPH both occurred in all 98 member proteins (**Figure 1b**). Both motifs were also highly conserved within Cluster 2, occurring in ≥90% of the 194 member proteins (**Figure 1c**). Although the GxSxG motif was used during identification of these lipases, the conserved version was identified to contain isoleucine and histidine as the variable second and forth positions, respectively. The type proteins of each lipase family/subfamily were scanned for the presence of the novel motifs to identify if they are unique markers of gut lipases. AHSKGG was identified as unique to lipase Cluster 1 and 2 while the TTxxTPH motif was identified within the type proteins for lipase family 1.2 and 1.5.

To better understand the variability within this collection of gut bacterial lipases, their features were assessed. The length of each protein was studied in relationship to its host of origin (**Figure 1d**). The mean length was 410 amino acids, ranging from 158 to 706. Interestingly, 40.5% of pig bacterial lipases were less than 300 amino acids long. However, out of the ten longest lipases, four originated from the pig gut, and the other six from humans. The pH preference of each lipase was predicted, with the majority being alkaline (**Figure 1e**). The percentage varied between hosts, with the murine gut having the highest percentage of alkaline lipases (94.9%) while the porcine gut had the lowest (69.6%).

The localisation of enzymes is highly important as secreted enzymes can diffuse away from the cell, while membrane-bound enzymes are limited to substrates within the vicinity of the cells. Therefore, the presence of signal peptides and transmembrane regions was determined. Most bacterial lipases (84.5%) were identified to not contain any known signalling peptide, indicating most are not secreted extracellularly (**Figure 1f**). Of those that did contain a signalling peptide, the most common was the Sec signal peptide (Sec/SPI), followed by the lipoprotein signal peptide (Sec/SPII). No enzymes were observed to contain the Tat signal peptide (Tat/SPI). The majority of lipases were observed to contain transmembrane regions, although the distribution was host-specific (60.8 - 88.8%) (**Figure 1g**). Up to seven transmembrane regions were predicted within a single lipase, although this was observed in only a single lipase from the human gut. The most common number of transmembrane regions was five (19.0 - 49.4%), followed by one (1.7 - 25.0%).

### Taxonomic and functional insights into lipase-positive bacteria

Metagenomic assembled genomes (MAGs) allow comprehensive insights into microbial diversity, including species that are currently uncultured [34–36]. Proteins from 154,723 MAGs previously reconstructed using metagenomes from different human body sites [36] were annotated using the same lipase annotation method as for the gene catalogues. In total, 13,677 lipases were identified within 11,011 of the 154,723 MAGs (7.1%). Most of these proteins were assigned to Cluster 2 (n = 7,604). However, many were also identified as being members of the sub-families within lipase family 1 (n = 1,487) [37]. Of those assigned to Cluster 2, 98.3% contained the three motifs (GxSxG, TTxxTPH and AHSKGG) along with 98.5% of the 653 proteins assigned to Cluster 1.

To further study the taxonomy of lipase-positive species, the MAGs were grouped based on average nucleotide identity (ANI) values, reducing the 154,723 MAGs into 4,930 species genome bins (SGB). For each SGB, a single representative MAG was selected, of which 235 (4.8%) were identified to contain a lipase (**Figure 2a**). Taxonomically, these lipase-positive species belong to 37 bacterial families across seven phyla: *Bacillota*, *Actinomycetota*, *Pseudomonadota*, *Fusobacteriota*, *Bacteroidota*, *Spirochaetota,* and *Cyanobacteriota*. Of the most prevalent SGBs (based on the number of MAGs assigned to the bin), only 8 of the top 23 could be assigned to a named species, whilst the rest represented metagenomically inferred species of novel lineages (**Figure 2b**). The mean relative abundance of lipase-positive species was nearly twice that of the lipase-negative species (1.06% vs 0.68%), suggesting that they are dominant members of human microbiomes (**Figure 2c**). Although dominant, almost 50% of the SGBs were identified to represent unknown species or genera according to GTDB-TK (**Figure 2d**), indicating that many lipase-positive species lack a cultured representative. The majority of the SGB representatives originated from gut samples (n = 176, 74.9%) and represented the major phyla from the gut (*Bacillota*, *Bacteroidota*, *Actinomycetota*, *Pseudomonadota,* and *Fusobacteriota*). The most commonly reconstructed lipase-positive species was *Ruminococcus_D bicirculans*, renamed to *Hominimerdicola aceti* [38], a dominant commensal of the human gut [39] which contains a lipase belonging to Cluster 2 (**Figure 2b**).

**Figure 2:**
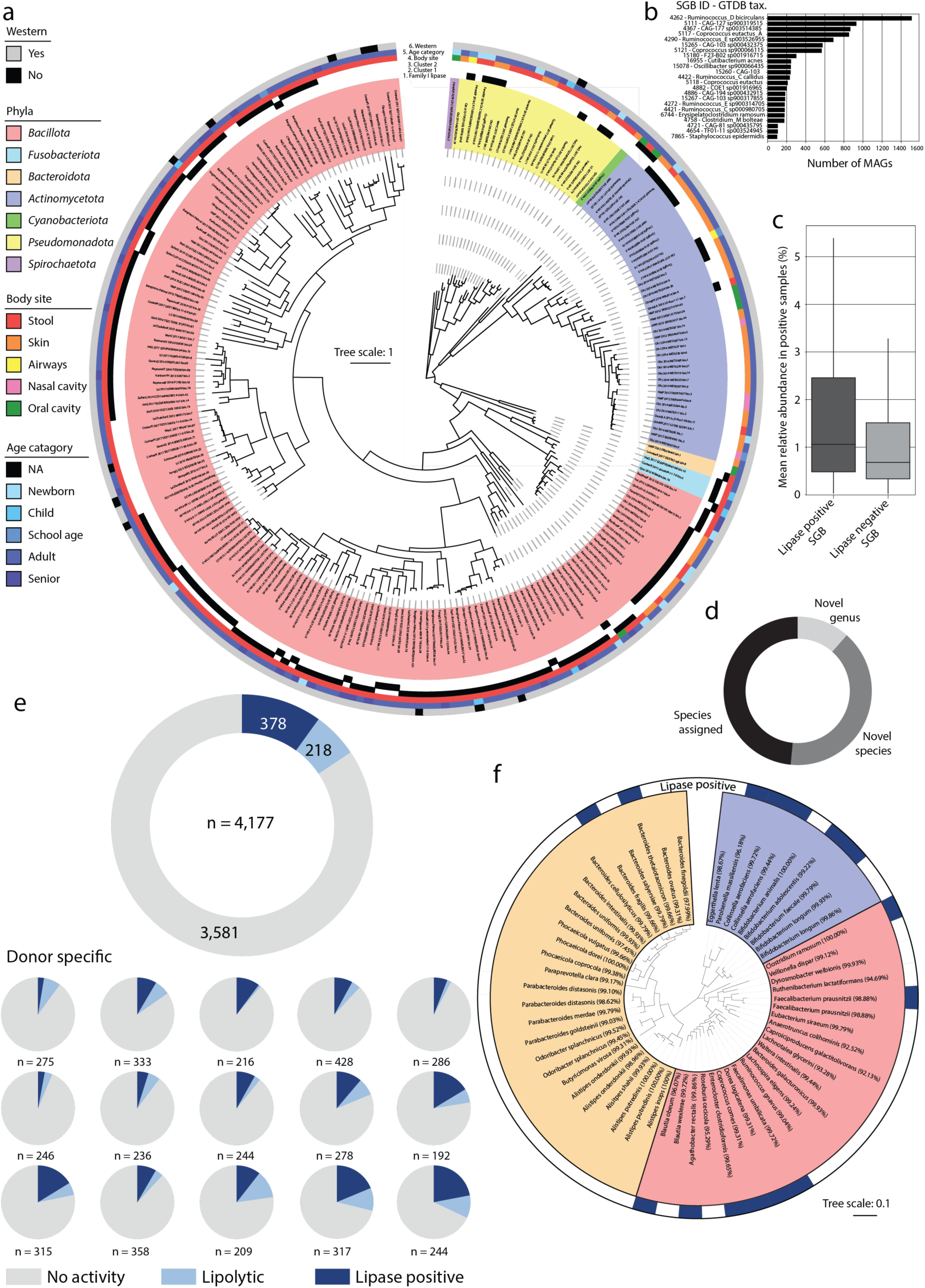
Culture-independent and -dependant diversity of lipase-positive species. **a.** Phylogenomic tree of 235 representative lipase-positive SGBs from human microbiomes generated using PhyloPhlAn [43]. External rings depict important features of SGBs as follows (inner to outer rings): lipase belonging to either family I, Cluster 1 or Cluster 2 (black, encoded; white, absent); metadata associated to the sample the SGB representative was reconstructed from: body site, age of the host, and if from a Western country or not (see colour legends). **b**. The taxonomy and size (number of MAGs) of all lipase-positive SGBs with a size > 100. **c**. Mean relative abundance of lipase-positive and -negative SGBs across 7,783 human gut metagenomes. **d**. Novelty of lipase-positive SGBs according to their GTDB taxonomy. **e.** Occurrence of lipolytic and lipase-positive bacteria across 4,177 isolates (top) or per donor (N = 15) (bottom). **f.** Phylogenetic tree (16S rRNA gene sequences) of a collection of 57 human gut isolates with their lipase activity, as determined by rhodamine B agar testing.

The second largest body site from which lipase-positive species were identified was the skin (45 SGB, 19.1%), of which the majority (29 SGB, 64.4%) belonged to the phylum *Actinomycetota* (**Figure 2a**). Additionally, nine *Bacillota* species belonging to the genus *Staphylococcus* were identified*. Staphylococcus* species have previously been shown to include lipase producers from the skin microbiota [40]. Both *Staphylococcus epidermidis* and *Cutibacterium acnes*, common skin colonisers and some of the most prevalent reconstructed species within the utilised dataset (**Figure 2b**), were identified to be lipase-positive, consistent with previous reports about these species [41].

In addition to the culture-independent study of MAGs, we determined the lipolytic activity of 4,177 bacterial colonies from faecal samples from 15 healthy human adults. Of these, 596 (13.5%) were assumed to express either a lipase or esterase based on screening using tributyrin agar (**Figure 2e**). Lipase activity was confirmed by screening the 596 isolates by re-streaking on rhodamine B agar. In total, 378 (8.6%) were lipase-positive, representing between 2.5 and 18% of the isolates from each donor. All donors were found to contain lipase-positive bacteria with a mean prevalence of lipase-positive colonies of 9.1% ± 4.6% (**Supplementary Table 1**), confirming lipase activity as a dominant phenotype.

To identify the taxonomic diversity of lipase-positive species in the human gut, a curated collection of 57 phylogenetically diverse strains representing 31 dominant genera was studied further [42]. Out of these strains, 15 were clearly identified to be lipase-positive based on the rhodamine B agar assay (**Figure 2f**). As this collection contained multiple strains for 6 species (>98.7% 16S rRNA gene sequence identity to the same assigned species), the consistency of lipase activity between strains could be assessed. While strains assigned to *Collinsella aerofaciens, Bifidobacterium longum*, *Alistipes putredinis*, and *Odoribacter splanchnicus* showed consistent results, strain-level variability was observed in *Faecalibacterium prausnitzii,* with one being positive and the other negative.

### Characterization of a novel bacterial lipase from the gut

The lack of experimental validation for bacterial functions has been highlighted as one of the major limitations of metagenomic studies [8]. Hence, we utilised cultured isolates and biochemical testing to provide *in vitro* evidence for their functional assignment. Genomes from in-house isolates [44] were searched against the predicted lipases to identify cultured host species. A complete sequence match (100% identity and coverage) was identified between a member of Cluster 2 and a protein within *Clostridium symbiosum* DSM 29356, a prevalent species in the human gut (detected in 28.10% (281/1,000) of human gut metagenomes) [45]. The isolate was cultured in a lipid-rich medium (see methods) for 24h and tested for lipolysis on rhodamine B agar. The fluorescence detected from the *C. symbiosum* cell solution, compared to the negative control (*Escherichia coli* DSM 28618) and the positive control (porcine lipase) confirmed its lipase activity (**Figure 3a**). To determine if this enzymatic activity was cell-bound or excreted, the cell pellet and supernatant were tested separately, with activity being identified only with the cell pellet (**Figure 3b**). Sequence analysis confirmed that the protein contains the GxSxG, AHSKGG, and TTxxTPH motifs, as well as five transmembrane regions at its N terminus, suggesting it is a membrane-bound lipase (**Figure 3c**). The predicted model of the lipase had a low normalised B-factor value for the predicted transmembrane region, indicating its flexibility and therefore supporting its prediction as a transmembrane region (**Figure 3d**).

**Figure 3:**
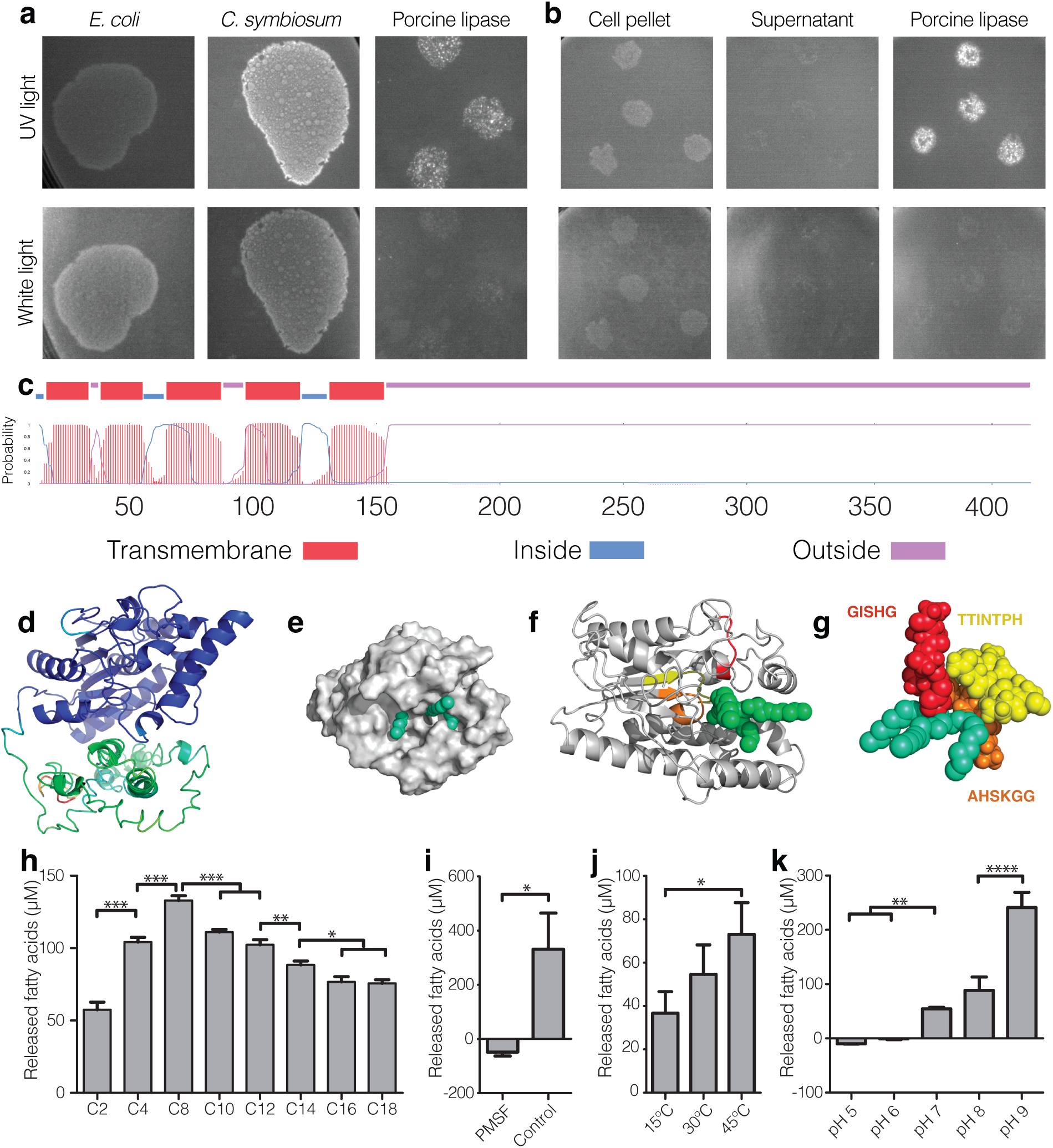
Structural modelling and biochemical characterisation of LipΔ150. **a.** Rhodamine B agar assay for *C. symbiosum* DSM 29356, *E. coli* DSM 28618 (negative control) and porcine lipase (positive control) viewed under both UV (60 ms) and white light (400 ms). Lipase activity is indicated by fluorescence under UV light, seen as white in the images for *C. symbiosum* and porcine lipase (top). **b**. Rhodamine B agar assay for the cell pellet and supernatant fractions of *C. symbiosum*, cultured for 7 days, along with the porcine lipase-positive control, viewed under both UV (top) and white light (bottom). **c**. Prediction of transmembrane regions via TMHMM [33]. **d**. Structural model of the complete lipase protein coloured according to the B-normalised value, indicative of structural flexibility, from green (low = - 1.16) to blue (high = 2.52). **e-g**. Structural modelling of LipΔ150 bound to the substrate OCP (green) on the surface (**e**), using a ribbon model (**f**) and highlighting only the conserved motifs (**g**). In both **f** and **g**, the conserved motifs are coloured red for GISHG, orange for AHSKGG, and yellow for TTINTPH. **h-k**. Biochemical characterisation of LipΔ150 was conducted using a range of fatty acid lengths (**h**), inhibition via PMSF (**i**) temperature-dependant activity (**j**), and pH (**k**). Triplicates were tested for statistical significance (t-test with Bonferroni multiple test correction) is represented as follows: * < 0.05, ** < 0.01, *** < 0.005.

Structural modelling predicted that removal of the transmembrane region at amino acid 150 of the amino acids sequence would leave a functional and stable enzymatic region (**Figure 3e-g**). Comparison of the predicted structure of the truncated lipase, LipΔ150, against the PDB database using COFACTOR confirmed that the five top matches, based on TM-score, were lipases. Functional annotation of the structure assigned the EC as 3.1.1.3 and the gene ontology term ‘triglyceride lipase activity (GO:0004806)’. Ligand binding was also predicted, based on a TM-SCORE of 0.81, of 5LipA bound to octyl-phosphinic acid 1,2-bis-octylcarbamoyloxy-ethyl ester (OCP), a structural homolog for triglycerides which binds irreversibly to the catalytic serine, facilitating crystallography (**Figure 3e**). The AHSKGG, TTxxTPH and GxSxG motifs were all identified to co-localise with OCP within the lipase structure (**Figure 3f**). Co-localisation of the motifs and OCP was studied further by space-filled modelling the motifs and OCP molecule (**Figure 3g**). The conserved nature of these motifs, along with their co-localisation with OCP suggests that they are not only hallmarks of gut lipase clusters but are of functional importance.

The LipΔ150 sequence was cloned with a poly-histidine tag into a pET-15b expression vector in *E. coli* BL21(DE3). Once expressed and purified, the enzyme was biochemically characterized to test its predicted function. LipΔ150 was confirmed to be a lipase, as activity was detected against all lengths of fatty acids tested, ranging from C2 to C18 (**Figure 3h**). However, a preference was shown for medium-chain fatty acids under the conditions tested, as the highest concentration of released fatty acids was observed with C8 (133 ± 3 µM), and the least was with C2 (57 ± 5 µM).

Due to the use of the GxSxG motif in the prediction of these enzymes, PMSF was used to identify if LipΔ150 is a serine-hydrolase. PMSF inhibited 100% of the lipase activity and led to mild precipitation of the protein (**Figure 3i**). Lipolytic activity was also observed to be temperature-dependent, with maximum activity occurring at 45 °C (**Figure 3j**). The possibility of pH altering lipolytic activity was also investigated, with pH of 6 or below preventing activity and maximum activity being observed at pH 9 (**Figure 3k**). As LipΔ150 represents a novel lipase based on its low sequence similarity (<40%) to all existing families, we propose it as the first member of the new lipolytic enzyme family XXXVI [30].

### Lipase-positive bacteria preferentially utilise glycerol rather than fatty acids

The presence of lipases within a substantial fraction of gut-colonizing-bacteria suggests a metabolic benefit to triglyceride degradation, such as for energy generation or biomass production [46]. To investigate this further, we examined the metabolic potential of the 235 lipase-positive SGBs (**Figure 4a**). Lipolytic activity releases two products: free fatty acids that can undergo beta-oxidation, and glycerol which can be metabolized to enter glycolysis. Three models of lipid catabolism were studied [47]. Whilst aerobic and anaerobic models of beta-oxidation were examined [48], none of the SGBs contained the anaerobic beta-oxidation pathway and 5.5% of lipase-positive SGBs contained the complete aerobic beta-oxidation pathway. Conversely, 41.7% of SGBs contained the glycerol degradation II pathway. Additionally, when genomes containing incomplete pathways were included, a further 28.1% of SGBs were identified to utilize glycerol.

**Figure 4:**
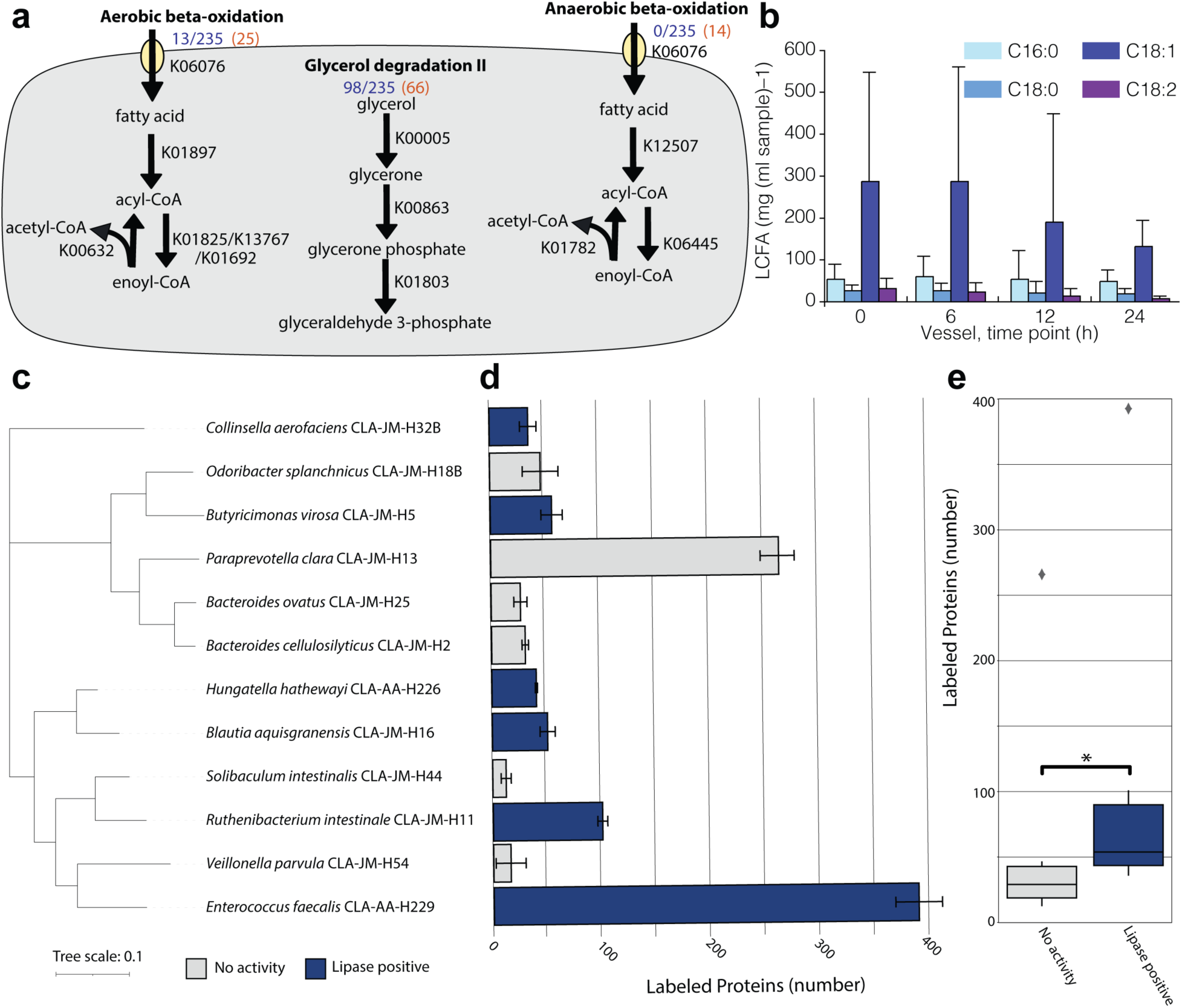
Glycerol utilisation by lipase-positive bacteria. **a.** The occurrence of each module for lipolytic catabolism across lipolytic SGBs is stated in blue; next to this, the number with the incomplete pathway (>50% of the total KOs in the pathway) is in orange. **b**. Degradation of long-chain fatty acids (LCFA) present in GMM and olive oil by faecal microbiota (n = 5) over 24 hours. The detected amount of each LCFA is plotted for each measured time-point during batch fermentation. Each microbiota originated from a different donor. **c**. Phylogenetic (16S rRNA gene sequence) tree of 12 bacterial strains isolated from the human gut, where 6 are lipase-positive (blue bars in panel c). **d**. Number of labelled proteins from each strain after incubation for 24 hours with C13-labelled glycerol. Bars represent the values across three replicates for each strain. **e**. Comparison of the mean number of labelled proteins for each strain between those that are lipase-positive, and those that are lipase-negative. Outliers are shown as grey diamonds. Statistical significance (Mann-Whitney U) is represented as follows: * < 0.05.

Due to the high-energy content of long-chain fatty acids, we initially assessed the ability of complex faecal communities to metabolise fatty acids (**Figure 4b**). For this, the faecal samples of five donors were incubated in GMM medium supplemented with 1g olive oil over 24 hours in duplicate. After 24 hours, the amount of C_18:1_ and C_18:2_ within the system was reduced from 149.2 ± 225.4 mg/ml to 69.2 ± 77.5 mg/ml (paired t-test, p=0.26) and 18.5 ± 21.0 mg/ml to 4.7 ± 4.6 mg/ml (paired t-test, p=0.06), respectively, although neither reduction was significant. The saturated fatty acids (C_16:0_, C_18:0_) remained at consistent levels throughout the 24 hours, suggesting the microbiota does not metabolise these substrates.

The genomic evidence suggests the microbiota produces lipases to access the glycerol subunits. To test this hypothesis, we studied the ability of six lipase-positive and six lipase-negative strains to utilise glycerol. We selected phylogenetically diverse species from both groups to ensure the phenotypes are not lineage-dependent (**Figure 4c**). To determine each strain’s ability to utilise glycerol, we incubated the strains in YCFA medium supplemented with 1% glycerol, of which 5% was C_13_ isotope labelled. Due to the isotope labelling, strains that utilised glycerol would have a fraction of their proteins labelled; hence proteomics coupled with protein stable isotope probing was used to determine the number of proteins labelled by each strain (**Figure 4d**). Interestingly, all strains were observed to contain labelled proteins, suggesting all could utilise glycerol to some extent. Comparison of the strains that were lipase-positive vs -negative strains indicated that the positive strains utilised glycerol to a greater extent, leading to a higher number of labelled proteins (**Figure 4e**).

## Discussion

The lack of detailed functional assignment is a limitation of metagenomic and genomic studies, limiting our insight into microbial processes. Therefore, we have generated a comprehensive collection of bacterial lipases present within the mammalian gut. The conserved yet unique sequences of these lipases, as compared to those isolated from other environments, was highlighted by the identification of two motifs which are both nearly ubiquitous across all identified bacterial lipases. We hypothesis that these motifs, along with the conserved GxSxG motif, are key to the activity of the bacterial lipases within the gut due to their co-localisation both together and with bound molecules. Of particular note is the high prevalence of alkaline lipases which corroborates previous findings that increasing the pH enhanced the activity of microbial phospholipases from the human gut [49].

Studying the taxonomic range of lipase-positive species identified that lipases are present across the major phyla and families within mammalian body-associated microbiomes. These results show lipolytic activity to be widespread, not limited to a few *Pseudomonadota* as previously suggested [47]. This discrepancy may be explained by the targeted annotation applied within this manuscript compared to the general annotations used to identify KEGG orthologs in previous studies [50].

Many of the most common lipase-positive SGBs were assigned to the genera *Ruminococcus* and *Coprococcus*, both of which are dominant members of the mammalian gut microbiome due to their diverse functional roles [51, 52]. Interestingly, the mean relative abundance of lipase-positive SGBs was twice that of lipase negative species, suggesting that these species are dominant and potentially key members of the ecosystem. The lack of aerobic or anaerobic beta-oxidation from the majority of lipase-positive SGB suggested that the metabolism of fatty acids is not the purpose of their lipolytic activity. In contrast, the majority of lipase-positive species were shown to utilise glycerol significantly more than strains without lipase activity, indicating the release and utilisation of glycerol may enhance the growth of lipase-positive microbes [53].

Direct measurement of lipolytic activity along the gastrointestinal tract can be difficult due to the diverse nature of intestinal content and sample variation. However, biochemical characterisation of a new gut bacterial lipase has provided some hints into their localisation and niche. Activity was observed to decrease with increasing detergent concentration, particularly above the critical micelle concentration. In addition to this, lipolytic activity was completely removed below pH 7, suggesting that bacterial lipases within the gut have evolved to work in an environment with sub-optimal detergent concentrations and at a pH at or above 7. Along the gastrointestinal tract, the bile concentration decreases whilst the pH and bacterial load increase, suggesting that the bacterial lipases have evolved to be active in the distal gut. For validation of this theory as well as to enhance our understanding of bacterial lipase localisation along the GI tract, *in vivo* experiments will be required.

## Materials and Methods

### Databases

#### EGGNOG

EGGNOG (v3) [54] consists of non-supervised orthologous gene clusters formed via Smith-Waterman reciprocal best matches and includes the original COGs [55], which have since been expanded. COG1075 is assigned the function ‘triacylglycerol lipase’ and contains 1,260 proteins from 633 species. The proteins originating from bacteria (67.5%) were used to form the EGGNOG database.

#### EC

Triglyceride lipases are assigned the enzyme commission (EC) notation 3.1.1.3, specifying them as first hydrolases, secondly acting on ester bonds, thirdly carboxylic ester hydrolases, and finally as triglyceride lipases. The UNIPROT database (accessed: December 2017) was searched for genes assigned this EC notation, which returned 162 proteins that formed the EC database.

#### Lipase engineering database (LED)

This database is a custom database of alpha/beta-hydrolase proteins encompassing six super families made up of 22 homologous families created from 5,278 proteins. Since its creation in 2000, the lipase engineering database has been used multiple times to study the sequence and structural similarity of hydrolases [29].

### Lipase annotation

Proteins from the pig [3], mouse [4] and human [2] metagenomic gene catalogues were annotated against the EC, EGGNOG and LED databases using DIAMOND BLASTP (v0.9.12.114) [56]. As previous research suggests protein families are best captured using 30-40% identity and 35-70% coverage [57], annotations were required to cover ≥40% of both the query and subject sequence at a minimum sequence identity of 40%. Moreover, proteins needed to have a match within two of the three databases and to contain the GxGxG motif identified to be common in lipases [29].

### Network analysis and multiple sequence alignment

Identity between the gut lipases and the type proteins for each known lipolytic protein family [30] was identified using DIAMOND with the same cutoffs as for lipase annotation. The protein matches were entered into Cytoscape (v3.8) [58] along with their shared identity for visualisation. The motifs specific for member sequences of both ‘Cluster 1’ and ‘Cluster 2’ (see results section) were identified via cluster-specific alignment with ClustalW [59] within the MEGAX software package [60] and visualised using weblogo [61].

### Protein modelling

The structure of protein sequences was modelled using the I-TASSER suite [62]. COFACTOR [63] was utilised with TM-ALIGN [64] to compare the predicted structure of the proteins to those within the Protein Data Bank (PDB) [65]. Models were viewed in Pymol (v1.8).

### Biochemical characterisation

The lipase sequence (WP_003509004.1) from *Clostridium symbiosum* DSM 29356 [44] was modified *in silico* by removal of the transmembrane region (first 150 N-terminal amino acids) and addition of a polyhistidine-tag (His_6_) at the C-terminus. The new construct (LipΔ150), integrated into a pET-15b vector was purchased from BioCat (Heidelberg, Germany) and transformed into *Escherichia coli* BL21 (DE3). The transformed *E. coli* was cultured in the presence of ampicillin (100 μg/ml) and a low IPTG concentration (0.1 mM) at low temperatures (25 °C) to support a functional folding of the expressed LipΔ150. Cells were harvested after 22 hours by centrifugation and frozen at -20 °C.

An aliquot of the frozen cells was solved in lysis buffer (50 mM Tris/HCl; 300 mM NaCl; 10 mM imidazole; pH 8) to a concentration of 150 mg/ml. The cells were disrupted by sonication cooled on ice (amplitude 25%; impulse length 3 x 30 s). Cell debris were removed by centrifuging the lysate (16,000 g, 30 min, 4 °C). The supernatant was filtered (0.45 µm) and the target protein purified via immobilized metal-ion affinity chromatography (IMAC) using a 1 ml HisTrap^TM^ HP column (GE Healthcare, UK). The lysis buffer (50 mM Tris/HCl; 300 mM NaCl; 10 mM imidazole; pH 8) was used for equilibration of the column and as washing buffer. Proteins were recovered using the elution buffer (50 mM Tris/HCl; 300 mM NaCl; 300 mM imidazole; pH 8) and collected in 0.5 ml fractions. The protein content during the washing and elution steps was monitored using the Äkta purifier 100 system (GE Healthcare, UK) and double-checked with a Bradford assay. Protein-containing fractions from the elution step were pooled and used for further purification.

Imidazole was removed using a PD-10 Desalting Column (GE Healthcare, UK) with an imidazole-free equilibration buffer (50 mM Tris/HCl; 300 mM NaCl; pH 8). Samples of approximately 200 µl were collected, a Bradford assay along with the *para*-nitrophenyl (*p*NP) assay (C8) (see below) were performed to identify the lipase fractions, which were then pooled. The final protein concentration was determined by a BSA-Bradford assay (R² 0.99). The purity of the isolated protein was verified with a native polyacrylamide gel electrophoresis.

Lipolytic activity was tested utilizing the degradation of *p*NP-bound fatty acid substrates [66–68]. The test was carried out by mixing 83.3 µl of a testing buffer with 16.7 µl protein solution and the absorption measured at 410 nm over a period of 45 min at 37 °C. The standard testing buffer (final concentrations in the 100 µl reaction mix: 50 mM Tris/HCl; 20 mM NaCl; 4.46 mM Tween20; 0.3 mM *p*NP-octanoate (C_8_); pH 8) was used during purification and for comparison in every test. A negative control was prepared for every tested condition by adding 16.7 µl of lipase free equilibration buffer (50 mM Tris/HCl; 300 mM NaCl; pH 8) to 83.3 µl of the corresponding testing buffer (*i.e.* containing *p*NP substrates of varying chain length). After 45 min incubation at 37 °C, samples (including all controls) were centrifuged (4,000 g, 10 min) and the supernatant measured. For quantification, a standard curve was created using the *p*NP assay with defined concentrations of *p*NP (R^2^ = 0.99). Results were normalized by subtraction of the negative control values and the standard curve was used to determine the concentration of released fatty acids. Statistical significance was determined using a t-test with Bonferroni multiple test correction adjusted p-values reported.

### Lipolytic activity assay of bacterial isolates

Hungate tubes with 9 ml of anaerobic brain–heart infusion (BHI) medium were used to cultivate *Escherichia coli* DSM 28618 (negative control) and *Clostridium symbiosum* DSM 29356. After 24 hours, 50 µl of each culture was placed onto rhodamine B agar plates [69]. To identify if the lipolytic activity was cell-bound, each isolate was cultured in Hungate tubes with 9 ml BHI broth supplemented with 0.5% cysteine (0.05 g/ml), 0.5% DTT (0.02 g/ml), and 200 µl emulsified olive oil. After 7 days of cultivation, 6 ml of dense culture was centrifuged (10,000 g, 10 min) to collect biomass. The pellet and 50 µl of the supernatant were put onto rhodamine B agar plates. In each experiment, 50 µl of porcine pancreas lipase (Sigma-Aldrich, USA) with a concentration of 20 mg/ml was added as positive control to every plate. Afterwards, the plates were incubated for 2 hours at 37 °C. Fluorescence, caused by lipolytic activity was observed under UV light using the GelStudio SA (Analytic Jena, Germany). Curated human gut isolates are all publicly accessible as members of the HiBC [42].

### Genome analysis of catabolism pathways

Metagenome-assembled genomes (MAGs) were obtained from Pasolli *et al.* (2019) [36], along with metadata for their originating sample, assignment to species genome bins (SGBs), prevalence and relative abundance [36]. Taxonomy was assigned to each MAG using GTDB-TK, release 89 [70, 71]. MAGs were annotated using PROKKA (v1.13.3) [72] and KEGG orthologs (KOs) [73] identified using the PROKKA2KEGG tool (https://github.com/SilentGene/Bio-py/tree/master/prokka2kegg). Each MAGs KOs were searched against previously published lipolytic catabolism modules [47]. Genome based trees were created using the predicted protein sequences in PhyloPhlAn (v3.0.60) [74] with the high diversity option.

### 16S rRNA gene sequence analysis

Annotation of 16S rRNA gene sequences was done using EZBioCloud [75]. The phylogenetic tree was generated using MegaX [76], with alignment conducted with MUSCLE [77] and a maximum likelihood tree generated using a GTR model with uniform rates of mutation and removal of missing data, bootstrapped via 1000 replicates. The tree was then annotated in iToL [78].

### Large-scale identification of lipase active strains from human faecal samples

Fresh faecal samples were obtained from 15 healthy adults (**Supplementary Table 1**). The collection of anonymised faecal samples used in this study has been described previously [79]. At the laboratory bench, samples were kneaded manually to make the samples homogenous. Sterile, anaerobic PBS (0.1 M, pH 7.2; Oxoid) was added to portions (∼5 g) of faeces in mini stomacher bags (Seward) to give 1:9 faeces-to-PBS (w/w). Samples were transferred to an anaerobic cabinet (Don Whitley; 80 % N_2_, 10 % H_2_, 10 % CO_2_) maintained at 37 °C. The total amount of time the faecal samples were exposed to air for was ∼3 min. Dilution series (10^−1^ to 10^−8^) were prepared from the faecal homogenates in sterile, anaerobic PBS.

Isolation was conducted on five agar media (fastidious anaerobe agar containing 5 % laked horse blood (Oxoid), reinforced clostridial agar (Oxoid), phenyl ethyl alcohol agar containing 5 % laked horse blood, Mueller– Rogosa–Sharpe agar (Oxoid) and Maconkey agar (Oxoid)). Plates were incubated for 5 days, then colonies were counted. Each colony from one of the replicates for each plate from which counts were obtained was picked and spotted onto tributyrin agar. Tributyrin agar was prepared by adding 5 g tributyrin agar (Sigma Aldrich) to 250 ml distilled water and 1.25 ml of glyceryl tributyrate (Sigma Aldrich). The mixture was autoclaved at 121 °C, 15 psi for 15 min. After autoclaving the molten agar was homogenized in a 250-ml Ato Mix blender (MSE) for 30 seconds. Rhodamine B agar was prepared in the same way as the tributyrin agar, except that the glyceryl tributyrate was substituted with olive oil and 2.5 ml of filter-sterilized rhodamine B (1 mg ml^−1^ solution prepared in distilled water) was added. Tributyrin plates were examined for zones of clearing at 4, 12, 24 and 48 hours. Colonies demonstrating lipolytic activity at any of these times were identified by marking the plates. Lipolytic bacteria were spotted onto the rhodamine B agar to confirm lipase activity as determined by zones of fluorescence detected under a UV light.

### Glycerol labelled proteomics

Biological triplicates of the selected bacterial strains were added to 10 ml of suitable liquid cultivation medium supplemented with 129.99 mM glycerol (Sigma Aldrich) and 6.84 mM ^13^C3-labelled glycerol (Eurisotop) (1% (v/v) glycerol of which 5% (v/v) are labelled) and incubated at 37 °C for 24 hours under anaerobic conditions. The cultures were centrifuged at 4,500 g for 10 minutes and the supernatant was removed. After washing the cells with 1 ml of PBS the cell pellets were immediately frozen and stored at -80 °C.

Bacterial pellets were resuspended in a lysis buffer consisting of 8 M urea, 2 M thiourea, and 1 mM phenylme-thylsulfonyl fluoride. To disrupt the bacterial cells, bead beating was performed using the FastPrep-24 system (MP Biomedicals, Santa Ana, CA, United States; 5.5 ms, 1 min, 3 cycles), followed by heat shock at 90 °C for 10 min with 1,400 rpm (ThermoMixerTM, Eppendorf, Germany) and centrifugation (10,000× g, 10 min). The resulting supernatant was used for protein concentration determination using a Pierce 660 nm Protein Assay (Thermo Fisher Scientific, Waltham, MA, United States). Vivacon 500 columns with a 10 kDA Molecular weight cutoff membrane (Sartorius, Göttingen, Germany) were equilibrated with lysis buffer and centrifuged (14,000× g, 20 °C, 20 min), and the remainder of the centrifugation steps were performed under the same conditions. Subsequently, 50 μg of the protein was loaded onto the column and centrifuged. The proteins that adhered to the filter were then incubated with 200 μL of 10 mM dithiothreitol in lysis buffer using a Thermo-Mixer (Eppendorf) at 37 °C and 600 rpm for 1 min and then with no agitation for 30 min. After centrifugation, the protein pellets underwent further alkylation and proteolytic cleavage, as described previously [80]. Finally, the resulting peptide lysates were dissolved in 0.1% formic acid (FA) for subsequent mass spectro-metric measurements.

For each nanoLC-MS measurement, 1 μg of the peptide lysate was injected into a Vanquish Neo nano-HPLC system (Thermo Fisher Scientific). The peptide lysates were initially trapped on a C18-reverse phase trapping column (Acclaim PepMapTM 100, 75 μm × 2 cm, particle size 3 μM, nanoViper, Thermo Fisher Scientific), and then separated on a C18-reverse phase analytical column (Double nanoViper™ PepMap™ Neo, 75 μm × 150 mm, particle size 2 μM, Thermo Fisher Scientific). The separation was achieved using a two-step gradient with mobile phases A (0.01% FA in H2O) and B (80% acetonitrile in H2O and 0.01% FA). During the first step of the gradient, which lasted 95 min, the proportion of mobile phase B was increased from 4 to 30%. This was followed by a 40-min period in which the proportion of mobile phase B increased from 30 to 55%. The flow rate during the separation was maintained at 300 nL/min as described [81]. The eluted peptides were ionized using a Nanospray Flex™ Ion Source (Thermo Fisher Scientific) and detected using an Orbitrap Exploris™ 480 mass spectrometer (Thermo Fisher Scientific). The mass spectrometer was operated with the following settings: for MS, the scan range was set to 350–1,550 m/z, the resolution was set to 120,000, the automatic gain control (AGC) target was set to 3′000,000 charges, and the maximum injection time was set to 100 ms. An intensity threshold of 8,000 ions and dynamic exclusion time of 20 s were also applied. The top 10 most intense ions were selected for MS/MS analysis, with an isolation window of 1.4 m/z, resolution of 15,000, AGC target of 200,000 ions, and a maximum injection time of 100 ms.

Proteins from each strain that incorporated labelled 13C after incubation for 24 hours with 13C-labelled glycerol were identified using Sipros (v4) [82]. The mass spectrometry raw files were extracted into FT1 and FT2 format files using Raxport (https://github.com/thepanlab/Raxport.net) with the mono (https://github.com/mono/mono) framework. The extracted FT2 files of the samples were searched against a target-decoy protein sequence database comprised of the protein sequences available for the strains on UniProtKB and sequences from MAGs for strains whose sequences were not available on UniProt, where the reverse sequences of these target proteins were implemented as decoys. Label-free searches were performed using samples from strains incubated without any 13C labels using Sipros (v4) against the target-decoy database, and all identified proteins from the unlabelled searches were subsequently used as references for the labelled searches with Sipros. The false discovery rate (FDR) of peptide identifications was filtered at 0.05 and 0.01 for the unlabelled and labelled searches, respectively, by adjusting the score threshold of PSMs. The search required at least two unique peptides for each protein/peptide group to be represented in the results. The isotopic abundance of proteins provided the values of the ‘average enrichment levels’. To confirm that the labelled peptides were not representing the natural abundance of 13C, we set the threshold of the average enrichment level to 1.07, the same value as that of the 13C natural abundance [83].

### Batch fermentation

Fresh faecal samples were obtained from five healthy adult donors of between 25 and 38 years of age (3 females, 2 males). None of the donors had taken antibiotics during the 6 months prior to the study, and none was taking probiotics or prebiotics. Freshly voided faecal samples were processed immediately by diluting them 1 in 10 (w/w) in pre-reduced sterile H_2_O and homogenizing them in a filter bag in a stomacher (Stomacher 400 Lab System; Seward) for 2 min at ‘high’ speed. For each donor, a batch culture vessel containing 150 ml of gut model medium (GMM) [84] was inoculated with 20 ml of the faecal homogenate and 30 ml of water containing 25 μl TWEEN^®^ 80 (Fisher Scientific) and 0 or 1 g of refined/virgin olive oil blend (Carbonell). The water containing the detergent and lipid was steamed for 10 min prior to being added to the batch culture vessel. GMM comprised (g l^−1^ distilled H_2_O unless stated otherwise; all reagents were purchased from Sigma Aldrich unless stated otherwise): starch from potato, 5.0; pectin from citrus fruit, 2.0; guar, 1.0; porcine gastric mucin (type III), 4.0; xylan from oatspelts, 2.0; arabinogalactan from larchwood, 2.0; inulin (Orafti), 1.0; casein from bovine salts, 3.0; peptone water (Oxoid), 5.0; tryptone (Oxoid), 5.0; bile salts no. 3 (Oxoid), 0.4; yeast extract (Oxoid), 4.5; FeSO_4_.7H_2_O, 0.005; NaCl, 4.5; KCl, 4.5; KH_2_PO_4_, 0.5; MgSO_4_.7H_2_O, 1.25; CaCl_2_.6H_2_O, 0.15; NaHCO_3_, 1.5; L-cysteine, 0.8; hemin, 0.05 g dissolved in 1 ml of 1 M NaOH. All components of the GMM were combined and autoclaved at 121 °C for 15 min. The final working volume of each batch culture vessel was 200 ml. The pH of each vessel (pH 6.8) was controlled automatically by the addition of 0.5 M HCl or 0.5 M NaOH. pH controllers were supplied by Electrolab. The contents of each vessel were stirred constantly. An anaerobic environment was maintained by constantly sparging the vessels with O_2_-free N_2_. The temperature of each vessel was maintained at 37 °C by use of a circulating waterbath connected to each fermentation vessel. Each set of experiments was run for 24 h.

Samples (5 ml) were withdrawn from each vessel at 0, 6, 12 and 24 h and immediately placed on ice. Aliquots (2× 2 ml) were centrifuged at 13,000 rpm (MSE MicroCentaur; Sanyo) for 10 min at room temperature. The supernatants were retained for analysis of LCFAs, and stored at -80 °C until processed.

### Quantification of LCFAs via gas chromatography

For each sample, the supernatants from 1 ml samples were thawed on ice and total fatty acids were extracted as described [85]. C_19:0_ internal standard (stock solution, 200 μg C_19:0_ / ml methanol) was used as the internal standard. Fatty acid methyl esters (FAMEs) were separated and quantified using a gas chromatograph (model 6890; Agilent Technologies UK Ltd, Stockport, UK) equipped with a flame-ionisation detector, quadrupole mass selective detector (model 5973N), injection port and a 50 m fused silica capillary column (internal diameter 0·25 mm) coated with a 0.2 mm film of cyanopropyl polysiloxane (CP-SIL 88; Varian Analytical Instruments, Walton-on-Thames, Surrey, UK). Total FAMEs in a 1 ml sample at a split ratio of 15:1 were determined using a temperature gradient programme (initial temperature 80 °C for 1 min; increased at a rate of 25 °C/min to 160 °C, which was held for 3 min; increased at a rate of 1 °C/min to 190 °C, maintained for 5 min; increased at a rate of 2 °C/min to 230 °C, held for 25 min). Helium was the carrier gas and the apparatus was operated at a constant pressure (20 psi) and flow rate (0·5 ml/min). Injector and detector temperatures were maintained at 250 and 275 °C, respectively. Peaks were identified by comparison of retention times with authentic FAME standards obtained from Sigma (Poole, Dorset, UK) and Matreya Inc. (Pleasant Gap, PA, USA).

## Acknowledgments

TCAH and TC conceived the study. TCAH conducted all bioinformatic analyses and supervised daily laboratory work. JM, TS, LH, DHG, and TF conducted wet lab experiments. TCAH, JM, LH, DHG, TS, and TC interpreted data. LE, LH, and TC provided guidance and access to essential materials. TCAH and TC secured funding and wrote the paper. All authors reviewed the manuscript and agreed with its final content.

TCAH was funded by an internal University Hospital RWTH Aachen START grant titled ‘KETOGUT’ as well as a RWTH Start-Up grant titled ‘ProtoBiome’. Access to the computational resources of the RWTH cluster were provided under grant; rwth2608. TC received funding from the German Research Foundation (DFG) Project-ID 403224013 – SFB 1382 and Project-ID CL 481/17-1 – SPP 2474. LH’s PhD, from which the cultivation (LH, DHG) and batch-culture work (LH) is derived, was funded by GlaxoSmithKline. R. John Wallace and Nest McKain (Rowett Institute of Nutrition and Health, Aberdeen) are thanked for hosting LH and training her in gas chromatography and analyses of data.

## Ethical approval and consent to participate

All donors were members of the laboratory who gave free consent for their samples to be used for the purposes of isolating and characterizing lipase-positive bacteria, and for molecular-based analyses and have since been destroyed. At the time the study was conducted (2006/2007) [86], the University of Reading’s Research Ethics Committee (UREC) did not require that specific ethical review and approval be given by UREC for the collection of faecal samples from healthy human volunteers to inoculate *in vitro* fermentation systems.

## Supplementary Tables

**Supplementary Table 1:** Large-scale screening of bacterial colonies from human faeces. For each donor five growth medium were used, and for each the total number of colonies, as well as those that were lipolytic and lipase-positive are stated.

